# Combining different CRISPR nucleases for simultaneous knock-in and base editing prevents translocations in multiplex-edited CAR T cells

**DOI:** 10.1101/2022.11.11.516008

**Authors:** Viktor Glaser, Christian Flugel, Jonas Kath, Weijie Du, Vanessa Drosdek, Clemens Franke, Maik Stein, Axel Pruß, Michael Schmueck-Henneresse, Hans-Dieter Volk, Petra Reinke, Dimitrios L. Wagner

## Abstract

Multiple genetic modifications may be required to develop potent off-the-shelf chimeric antigen receptor (CAR) T cell therapies. Conventional CRISPR-Cas nucleases install sequence-specific DNA double-strand breaks (DSBs), enabling gene knock-out (KO) or targeted transgene knock-in (KI). However, simultaneous DSBs provoke a high rate of genomic rearrangements which may impede the safety of the edited cells. Here, we combine a non-viral CRISPR-Cas9 nuclease-assisted KI and Cas9-derived base editing technology for DSB free KOs within a single intervention. We demonstrate efficient insertion of a CAR into the T cell receptor alpha constant (TRAC) gene, along with two KOs that silence major histocompatibility complexes (MHC) class I and II expression. This approach reduced translocations to 1.5% of edited cells. Small insertions and deletion at the base editing target sites indicated guide RNA exchange between the editors. This was overcome by using CRISPR enzymes of distinct evolutionary origins. Combining Cas12a Ultra for CAR KI and a Cas9-derived base editor enabled the efficient generation of triple-edited CAR T cells with a translocation frequency comparable to unedited T cells. Resulting T cell receptor- (TCR-) and MHC-negative CAR T cells resisted allogeneic T cell targeting in vitro. Thus, we demonstrate a solution for safer multiplex-edited cell products and a path towards off-the-shelf CAR therapeutics.

## II. Introduction

Gene editing has become a central technology to engineer improved and more accessible cellular therapies to treat chronic diseases with high unmet medical needs, such as cancer or autoimmune diseases^1–3^. Programmable nucleases, such as Zinc-finger nucleases^4^, TALE nucleases^5^ or CRISPR-Cas^6,7^, enable the induction of DNA double-strand breaks (DSBs) at precise locations in the genome. Repetitive DSBs can be exploited for mutagenesis by provoking insertions and deletions (indels) via error-prone non-homologous end joining (NHEJ). Co-delivery of a programmable nuclease and a homologous DNA repair template can be used to install new genetic sequences at a precise location via homology-directed repair (HDR)^8^. Further, advanced gene editing enzymes are being developed that allow programmable changes of distinct bases^9,10^, epigenetic states^11–13^, or larger sequence changes, diversifying our toolbox to engineer the genome.

Chimeric antigen receptor (CAR) reprogrammed T cells are an increasingly important treatment modality for advanced cancers. The approved CAR T cell products rely on personalized manufacturing and retroviral gene transfer. Allogeneic CAR T cell therapy promises lower costs per treatment dose, and it could avoid treatment delays associated with autologous cell manufacturing. A universal T cell therapy from healthy human donors must overcome multiple hurdles: The T cells must (1) be effectively reprogrammed towards the cancer via a CAR (or transgenic TCR), (2) avoid GvHD caused by allo-reactive endogenous donor TCRs^14^, and (3) be protected from immediate allo-rejection by the host’s immune system^15^. These challenges can be overcome by modifying multiple genes at once (multiplex-editing)^1,16^.

By design, multiplex editing using CRISPR/Cas nucleases induces multiple DNA DSBs, which can induce chromosomal translocations due to the employment of DNA repair mechanisms without proof-reading function. Translocations can result in gene fusions or altered gene regulation, which have been associated with carcinogenesis^17^. A prominent example of gene fusion is the so called Philadelphia chromosome that encodes a hybrid BCR-ABL1 protein leading to CML^18^. Altered gene regulation has been observed in T cell malignancies where translocation of regulatory elements of the TCR genes modified expression patterns at other genomic sites, triggering malignant transformation^19^. Cell line experiments have shown that clones carrying driver mutations for cancer exhibit a reduced sensitivity for DNA damage. This can lead to selection and enrichment of these clones after a population has been subject to CRISPR-induced DNA DSBs^20^. Therefore, mitigating translocations and preventing DNA damage may reduce risks associated with multiplex editing of cellular therapies.

Previous T cell studies demonstrated that translocations decrease cellular fitness, leading to reduced expansion *in vitro*^21,22^ and in patients^23,24^. Recently, a clinical trial with multiplex-TALEN-edited CAR T cells was halted due to the detection of CAR T cells bearing chromosomal translocations in a patient with severe hematological toxicity (#NCT04416984; clinicaltrials.gov). The trial was continued, because the chromosomal abnormality was neither caused by TALEN gene editing nor was it found in the edited product^25^. This highlights that translocations remain a concern in the clinical translation of multiplex gene-edited T cell products and should be avoided.

Base editing combines the programmability of the CRISPR-Cas system with deaminase enzymes^9,10^. By fusing single strand DNA deaminase enzymes to a Cas9 nickase or an enzymatically dead Cas9 protein, targeted base changes can be introduced within a target window without a DNA DSB. This can be used to introduce stop codons or edit splice sites and consequently disrupt gene expression in T cells^26,27^. Furthermore, multiplex base editing has been combined with viral gene transfer to create potent CAR T cell products ^28,22,26^. So far, base editing has not been combined with CAR or TCR knock-ins (KI) to create redirected multiplex-edited T cells in a single gene editing procedure.

Here, we demonstrate a non-viral method that allows efficient KI of a therapeutically relevant transgene and silencing of additional genes after a single manipulation. We explore different CRISPR-Cas gene editors and characterize the rate of translocations and editing outcomes at the targeted sites by flow cytometry, digital droplet (dd)PCR and targeted next-generation sequencing (NGS). As a proof of principle, we used nuclease-assisted insertion of a CAR into the *T cell receptor alpha constant (TRAC)* locus and combined it with knockout (KO) of the *beta-2 microglobulin (B2M)*^29^ and *class II major histocompatibility complex transactivator (CIITA)* ^30,31^genes to abrogate surface expression of MHC class I and II molecules, respectively. Loss of MHC class I and II protects from allo-specific CD8 and CD4 T cell responses^16,32–34^ and does not impede the anti-tumor function of allogeneic CAR T cells *in vivo*^34^. The resulting TCR- and MHC-negative CAR T cells may represent a potential product candidate for off-the-shelf applications.

## III. Results

### HDR-mediated KI and double KO induces high rate of translocations

For proof-of-principle, we aimed to combine nuclease-mediated in frame-insertion of a 2^nd^ generation CD19-specific CAR transgene into the *TRAC* locus for functional TCR-to-CAR replacement^35–37^. Further, in the same intervention, we performed a double KO of *B2M* and *CIITA* to abrogate surface expression of MHC I and II, respectively (**Figure 1a**). DNA breaks at the three genes simultaneously could lead to genomic translocations potentially resulting in >30 unique chromosomal rearrangements (Fehler! Verweisquelle konnte nicht gefunden werden.**b**). Unbalanced translocations are greatly reduced during T cell expansion, while balanced translocations can persist in T cells^21^. Therefore, we established ddPCR assays to quantify the six balanced translocations between *TRAC, B2M* and *CIITA* as an estimate for the rate of translocations in our subsequent experiments (**Figure 1c**).

**Figure 1:**
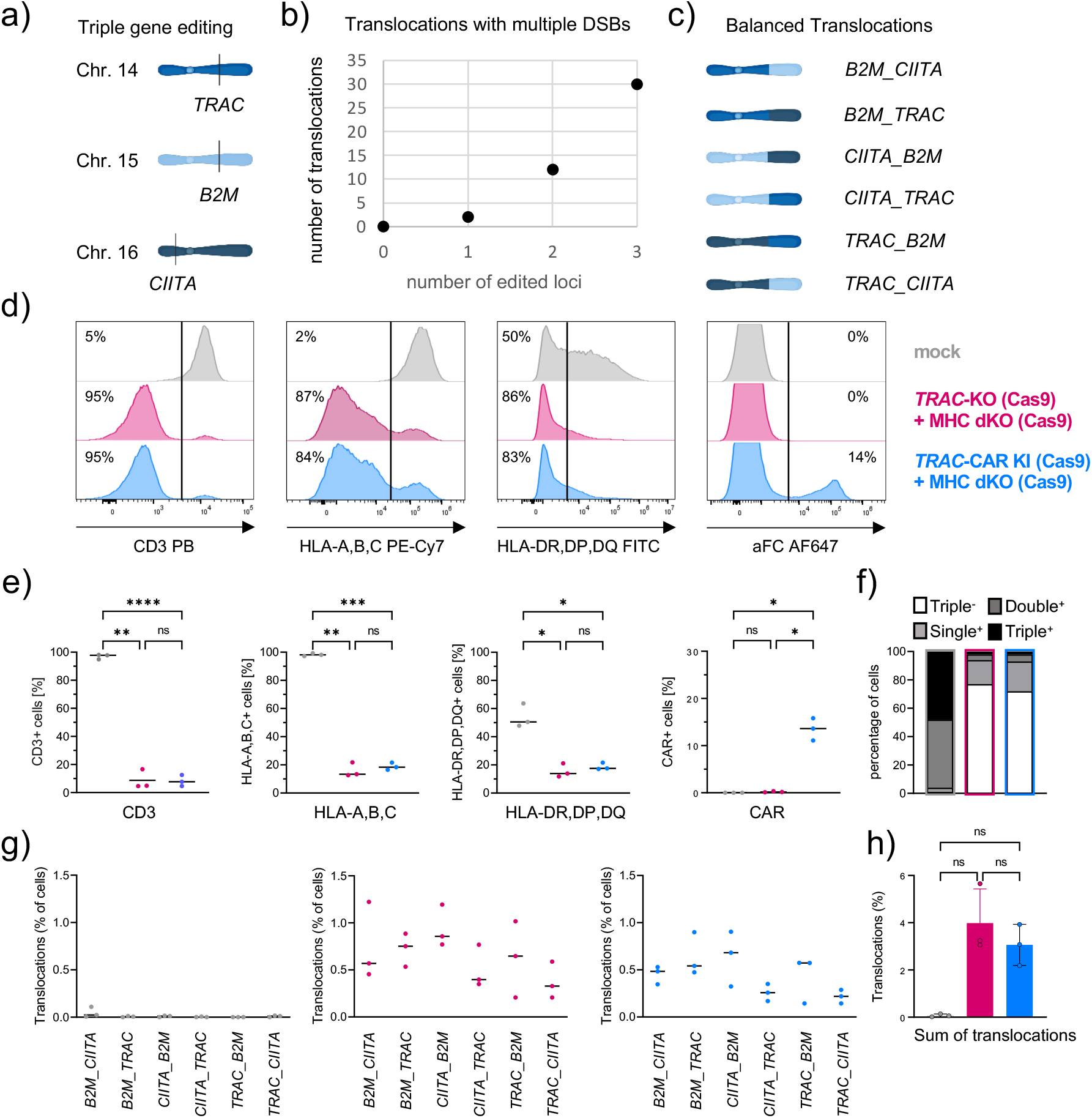
Conventional Cas9-mediated KI and double KO induces high rate of translocations. a) Triple gene editing strategy to generate allo-CAR T cells by harnessing CRISPR-Cas to target the *TRAC, B2M* and *CIITA* locus, leading to the depletion of the TCR as well as MHC class-I and –II molecules, respectively. b) Increase of unique translocations with increasing number of introduced double strand breaks (DSBs) 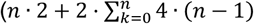 with n=number of introduced DSBs). c) Visualization of the six unique balanced translocations between the three targeted loci. d) Representative flow cytometry histograms show editing outcomes four days after co-transfection of Cas9 RNPs targeting the *TRAC, B2M* and *CIITA* genes alone (*TRAC*-KO (Cas9) + MHC dKO (Cas9)) or in combination with a homology directed repair template (HDRT) that facilitates the insertion of a CD19 transgene (*TRAC*-CAR KI (Cas9) + MHC dKO (Cas9)) in comparison to mock electroporated cells. The CAR was stained by using an aFC antibody targeting the IgG1 hinge. e) Summary of flow cytometry data of 3 individual healthy donors (n=3 healthy donors). f) Percentage of cells that are triple negative or positive for one, two or all three analyzed surface markers as determined by applying flow cytometry based on Boolean gating. g) Frequencies of cells carrying balanced translocations as determined by ddPCR are shown for all six individual translocations and h) as the sum of all translocations detected in mock, *TRAC*-KO (Cas9) + MHC dKO (Cas9)and *TRAC*-CAR KI (Cas9) + MHC dKO (Cas9) samples from n=3 donors. Statistical analysis of flow cytometry and ddPCR data from 3 donors was performed using a one-way ANOVA of matched data with Geisser-Greenhouse correction. Multiple comparisons were performed by comparing the mean of each column with the mean of every other column and corrected by the Turkey test. Asterisks represent different p-values calculated in the respective statistical tests (ns : p ≥ 0.05; * : p < 0.05; ** : p < 0.01; *** : p < 0.001; **** : p < 0.0001)

As a positive control to detect translocations, we transfected *Streptococcus pyogenes* (Sp)Cas9 protein and sgRNA targeting *TRAC, B2M* and *CIITA* to engineer triple KO cells (*TRAC*-KO (Cas9) + MHC dKO (Cas9)) using nucleofection. To create TCR-replaced CAR T cells, we co-electroporated a double-strand (ds)DNA homology-directed repair template (HDRT) to insert the CAR into the *TRAC* locus (*TRAC*-CAR KI (Cas9) + MHC dKO (Cas9)). As expected, the expression of CD3, HLA-A,B,C (MHC class I) and HLA-DR,DP,DQ (MHC class II) was significantly reduced four days after editing T cells in both *TRAC*-KO (Cas9) + MHC dKO (Cas9) and *TRAC*-CAR KI (Cas9) + MHC dKO (Cas9) (Fehler! Verweisquelle konnte nicht gefunden werden.**d,e**). CAR expression was observed in 13.5% ± 2.4% (mean ± SD, n=3) (**Figure 1e**). 77% of tKO and 72% of *TRAC*-CAR KI (Cas9) + MHC dKO (Cas9) were negative for all three markers (Fehler! Verweisquelle konnte nicht gefunden werden.**f**). All six balanced translocations were found in *TRAC*-KO (Cas9) + MHC dKO (Cas9) and *TRAC*-CAR KI (Cas9) + MHC dKO (Cas9) T cells (**Figure 1g**). In total, 4.1% ±1.2% (mean ± SD, n=3) of *TRAC*-KO (Cas9) + MHC dKO (Cas9) T cells displayed at least 1 balanced translocation (**Figure 1h**). Co-transfection of the HDRT in *TRAC*-CAR KI (Cas9) + MHC dKO (Cas9) reduced the rate to 3.4% ± 0.8% (mean ± SD, n=3), although this difference was not statistically significant (**Figure 1h**).

### Simultaneous Cas9 based KI and base editing reduces translocations between edited loci

Base editing allows the efficient disruption of multiple genes in T cells without causing translocations^26^. Therefore, we tested whether the Cas9 mediated KI could be combined with base editing to mitigate chromosomal rearrangements. To this end, we generated *in-vitro* transcribed, base modified mRNA of the adenine base editor ABE8.20m as previously described^38^. In pilot experiments, transfection of the ABE mRNA and single guide RNA (sgRNA) targeting a splice site of *B2M* or *CIITA*, we observed almost complete target base conversions in T cells highlighting high efficacy with the in-house produced mRNA as detected by Sanger sequencing and targeted NGS (Fehler! Verweisquelle konnte nicht gefunden werden.).

When combining Cas9 KI with the base editor (BE), we aimed to reduce gRNA exchange by pre-complexing the *TRAC* guide RNA (gRNA) with the SpCas9 protein and HDRT. Just immediately prior to electroporation, mRNA and gRNAs targeting *B2M* and *CIITA* were mixed with the Cas9 ribonucleinprotein (RNP) complex, the HDRT and the cell suspension (*TRAC*-CAR KI (Cas9) + MHC dKO (BE)) (**Figure 2a**). In comparison to *TRAC*-CAR KI (Cas9) + MHC dKO (Cas9), we observed similar silencing efficacy of CD3, HLA class I and class II molecules and comparable CAR integration rates in *TRAC*-CAR KI (Cas9) + MHC dKO (BE) treated T cells (Fehler! Verweisquelle konnte nicht gefunden werden. **b,c,d**). Further, we detected a 56% decrease in translocations from 3.4 ± 0.8% (mean ± SD, n=3) to 1.5 %± 0.5% (mean ± SD, n=3) relative to *TRAC*-CAR KI (Cas9) + MHC dKO (Cas9) (Fehler! Verweisquelle konnte nicht gefunden werden. **e,f**). Notably, still more than 1 in 100 cells harbored a balanced translocation between *B2M, CIITA* or *TRAC* overall (Fehler! Verweisquelle konnte nicht gefunden werden.**f**). These results indicate that, despite pre-complexation of the *TRAC* gRNA with the SpCas9 nuclease, the gRNA can be exchanged between the SpCas9 nuclease and the SpCas9-based BE provoking undesired DNA DSBs in *B2M* and *CIITA*.

**Figure 2:**
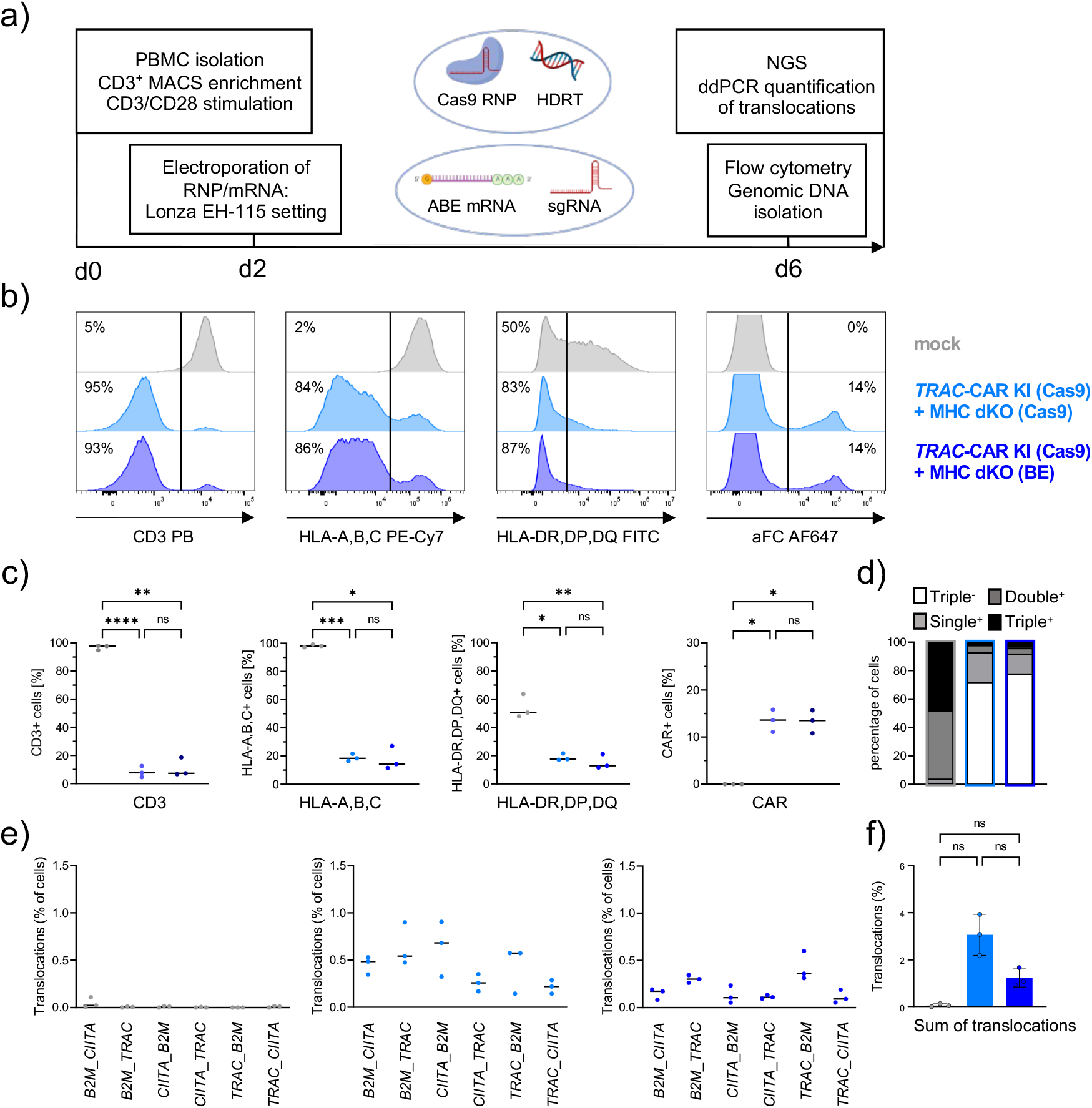
Co-delivery of Cas9-derived BE reduces translocations during simultaneous KI. a) Experimental setup for the generation of triple edited CAR T cells by co-delivery of a Cas9 RNP mediating the *TRAC* insertion with sgRNAs directing an mRNA encoded adenine base editor to target splice sites of the *B2M* and *CIITA* loci (*TRAC*-CAR KI (Cas9) + MHC dKO (BE)). b) Representative flow cytometry histograms show editing outcomes of *TRAC*-CAR KI (Cas9) + MHC dKO (Cas9), *TRAC*-CAR KI (Cas9) + MHC dKO (BE) and mock electroporated cells. c) Summary plots for surface expression data from 3 donors. The CAR was stained by using an aFC antibody targeting the IgG1 hinge. d) Percentage of cells that are triple negative or positive for one, two or all three analyzed surface markers as determined by applying flow cytometry based Boolean gating. e) Frequencies of cells carrying balanced translocations as determined by ddPCR are shown for all six individual translocations and f) as the sum of all translocation detected in mock, *TRAC*-CAR KI (Cas9) + MHC dKO (Cas9) and *TRAC*-CAR KI (Cas9) + MHC dKO (BE)samples from n=3 donors. Statistical analysis of flow cytometry and ddPCR data from 3 donors was performed using a one-way ANOVA of matched data with Geisser-Greenhouse correction. Multiple comparisons were performed by comparing the mean of each column with the mean of every other column and corrected by the Turkey test. Asterisks represent different p-values calculated in the respective statistical tests (ns : p ≥ 0.05; * : p < 0.05; ** : p < 0.01; *** : p < 0.001; **** : p < 0.0001)

### CAR KI using the Cas12a Ultra enzyme can be enhanced by mutating the PAM on the repair template

Combining different Cas enzymes should prevent gRNA exchange, as different nuclease proteins require different gRNA scaffold sequences for assembly of the RNP complex (**Figure 3**). Therefore, we hypothesized that CAR KI with the engineered Cas12a Ultra nuclease^39^ (derived from *Acidaminococcus sp*. Cas12a^40^) should allow for translocation-free gene editing when co-delivered with the SpCas9-based BE. The previously used HDRT for TCR-to-CAR replacement contained the target site for our *TRAC*-specific crRNA including a suitable PAM sequence. This means that the template could be cleaved and cause a reduced KI efficiency (**Figure 4a**). Surprisingly, the original HDRT yielded comparable CAR KI rates to Cas9 KI (**Figure 4 b,c**). Mutating the Cas12a PAM in the right homology arm of the HDRT resulted in a significant 2.9-fold (SD=0.9) increase of HDR rates with Cas12a averaging almost 30% KI (SD=9.5%) (**Figure 4d,e**). The new PAM-mutated HDRT did not increase the KI rate with Cas9 (**Figure 4f**). Overall, non-viral Cas12a KI with the PAM-mutated HDRT was 2-fold (SD=0.5%) more efficient than Cas9-KI with the same HDRT and parallel execution (**Figure 4g**).

**Figure 3:**
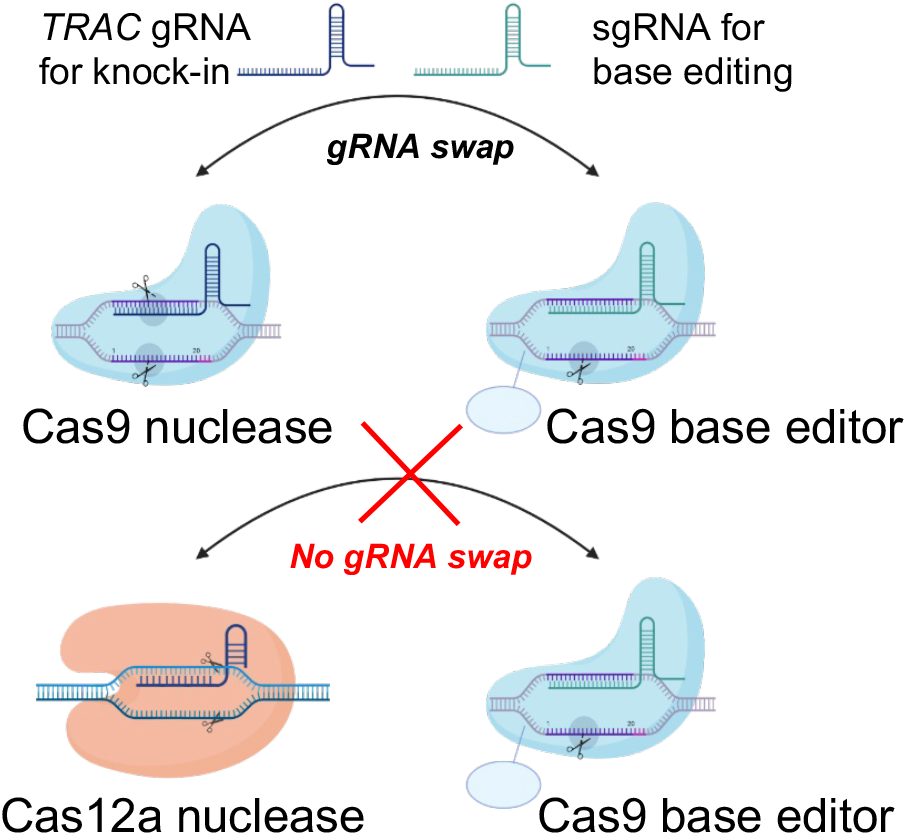
Guide RNA (gRNA) exchange can be avoided by using different Cas species for KI and base editing. Graphical display of gRNA exchange between the Cas9 nuclease and the Cas9 BE that can be prevented by applying a Cas12a nuclease in combination with a Cas9 base editor.

**Figure 4:**
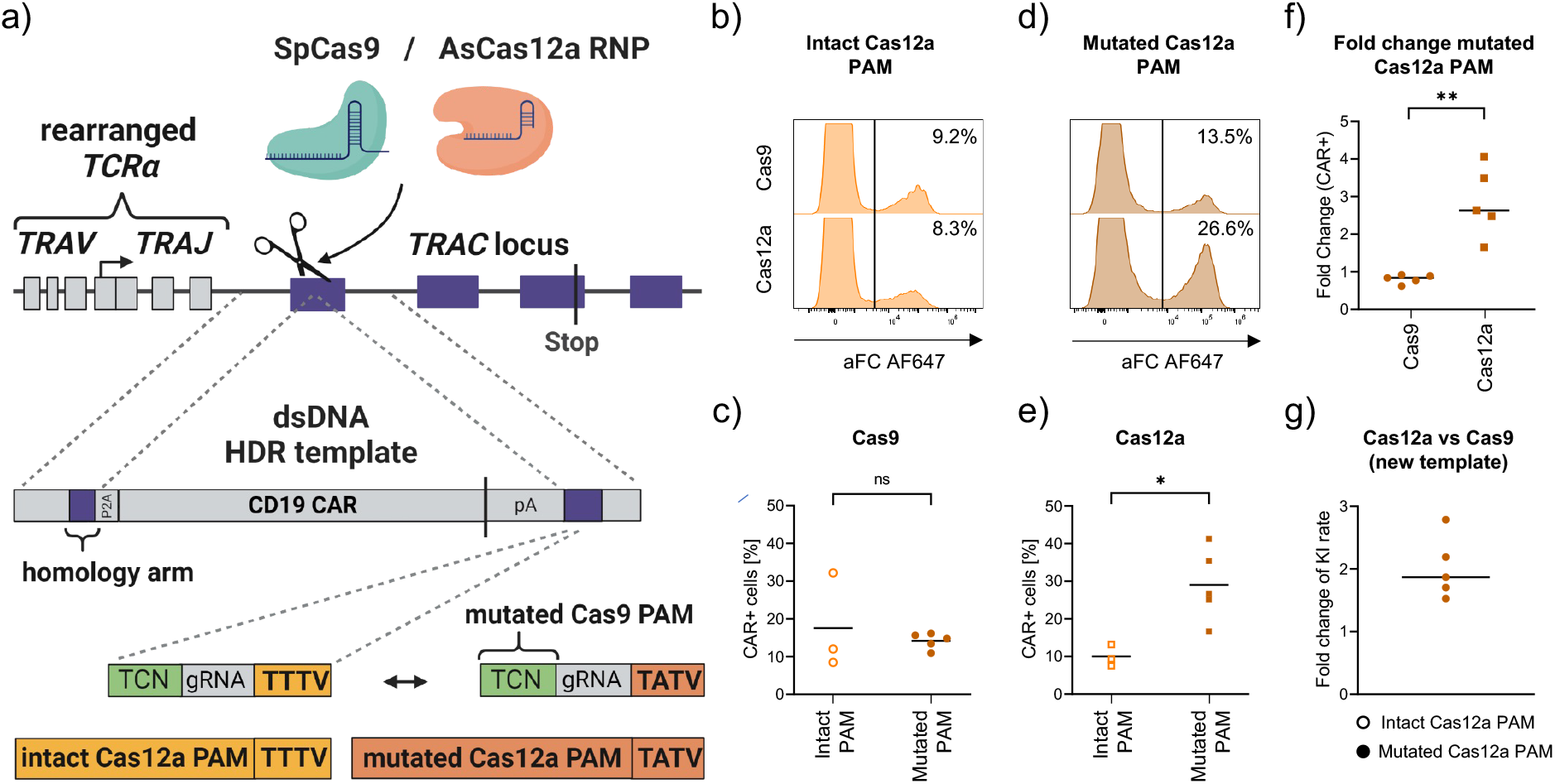
Eliminating Cas12a PAM in the HDR template increases KI efficacy. a) Design of the dsDNA template for *TRAC* targeted insertion is shown. The original template containing a mutated Cas9 PAM but an intact Cas12a PAM was modified by mutating the Cas12a PAM on the right homology arm to improve HDR efficiency by preventing cleavage of the repair template. b) Representative flow cytometry histograms show KI-efficiency using a template with an intact Cas12a PAM and either a Cas9 or Cas12a nuclease to target the *TRAC* gene. The CAR was stained by using an aFC antibody targeting the IgG1 hinge. c) Knock in efficiency quantified by flow cytometry using the intact or mutated template with a Cas9 nuclease (intact: n=3, mutated: n=5 individual donors; unpaired t-test). Empty shapes were performed with the old HDRT, filled shapes were performed with PAM-mutated HDRT d) Representative flow cytometry histograms show KI-efficiency using a template with the mutated Cas12a PAM and either a Cas9 or Cas12a nuclease to target the *TRAC* gene. e) Knock in efficiency using the intact or mutated template with a Cas12a nuclease (intact: n=3, mutated: n=5 individual donors; unpaired t-test). f) Fold change of KI efficiency with a donor template containing the mutated Cas12a PAM (n=5) in comparison to the mean (n=3) of the KI efficiency with the intact template (paired t-test). g) Fold change increase of KI efficiency by using Cas12a instead of Cas9 as a nuclease with the HDRT with mutated PAM (n=5). Asterisks represent different p-values calculated in the respective statistical tests (ns : p ≥ 0.05; * : p < 0.05; ** : p < 0.01; *** : p < 0.001; **** : p < 0.0001)

### Combining the Cas12a Ultra nuclease for KI and SpCas9 BE for KO allows efficient multiplex-editing and prevents translocations in T cells

When combining Cas12a KI and SpCas9-derived BE (*TRAC*-CAR KI (Cas12a) + MHC dKO (BE)) (**Figure 5a**), BE-mediated silencing of MHC class I and II was as efficient as in *TRAC*-CAR KI (Cas9) + MHC dKO (BE) (**Figure 5b,c**). Overall, >80% of T cells were negative for CD3, MHC class I and II after a single *TRAC*-CAR KI (Cas12a) + MHC dKO (BE) intervention (**Figure 5d**). The median fluorescent intensity (MFI) of MHC class I measured by HLA-A,B,C surface expression was significantly higher in *TRAC*-CAR KI (Cas12a) + MHC dKO (BE) than in *TRAC*-KO (Cas9) + MHC dKO (Cas9) (**Figure 5b, Suppl. Fig. 2d)**. Of note, co-electroporation of BE mRNA during Cas12a KI did not significantly impact the CAR T cell viability or expansion *in vitro* (**Suppl. Fig. 3**). Importantly, in *TRAC*-CAR KI (Cas12a) + MHC dKO (BE) treated cells, the translocations were significantly reduced by 10-fold to 0.14% (SD=0.07) (Fehler! Verweisquelle konnte nicht gefunden werden. **e,f**). Therefore, combining Cas12a Ultra mediated KI with Cas9-based BE enables complex genome engineering at multiple loci with minimal translocation.

**Figure 5:**
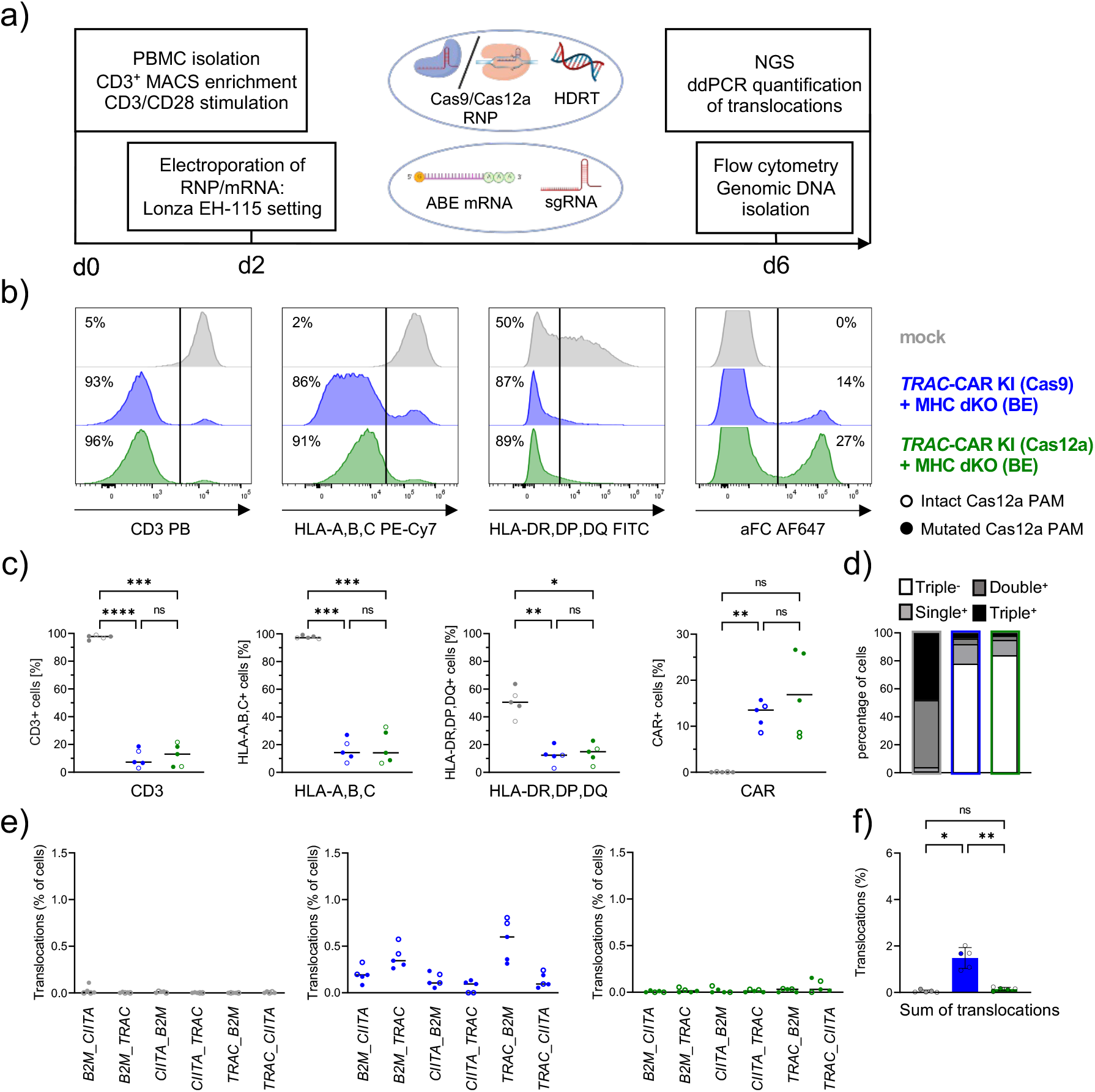
Co-delivery of Cas12a for KI and Cas9-derived BE avoids translocations during complex editing. a) Experimental setup for the generation of CAR T cells by co-delivery of a Cas9 or Cas12a RNP mediating the *TRAC* insertion with sgRNAs directing an mRNA encoded ABE to target splice sites of the *B2M* and *CIITA* loci (*TRAC*-CAR KI (Cas9) + MHC dKO (BE), *TRAC*-CAR KI (Cas12a) + MHC dKO (BE)). b) Representative flow cytometry histograms show editing outcomes of *TRAC*-CAR KI (Cas9) + MHC dKO (BE), *TRAC*-CAR KI (Cas12a) + MHC dKO (BE) and mock electroporated cells. The CAR was stained by using an aFC antibody targeting the IgG1 hinge. c) Summary plots for surface expression data from 5 donors. Empty shapes were performed with the old HDRT, filled shapes were performed with PAM-mutated HDRT. d) Percentage of cells that are triple negative or positive for one, two or all three analyzed surface markers as determined by applying flow cytometry based Boolean gating. e) Frequencies of cells carrying balanced translocations as determined by ddPCR are shown for all six individual translocations and f) as the sum of all translocations detected in mock, *TRAC*-CAR KI (Cas9) + MHC dKO (BE) and *TRAC*-CAR KI (Cas12a) + MHC dKO (BE) samples from n=5 donors. Statistical analysis of flow cytometry and ddPCR data from 5 donors was performed using a one-way ANOVA of matched data with Geisser-Greenhouse correction. Multiple comparisons were performed by comparing the mean of each column with the mean of every other column and corrected by the Turkey test. Asterisks represent different p-values calculated in the respective statistical tests (ns : p ≥ 0.05; * : p < 0.05; ** : p < 0.01; *** : p < 0.001; **** : p < 0.0001)

### Next-generation sequencing confirms lack of indels at Cas9-BE targets when using Cas12a for KI

Adenine base editing should induce targeted A to G conversions without indels. In contrast, gRNA exchange between the SpCas9 nuclease and the BE may cause undesired DSBs at BE-target sites, leading to indels (**Figure 6 a,b, Suppl. Figure 4**). Amplicon sequencing of the different gene-edited CAR T cell products revealed high rates of indel formations in *TRAC*-CAR KI (Cas9) + MHC dKO (BE) (*B2M*: 54.8% ± 5.7%, *CIITA*: 19.4% ± 3.1%, mean ± SD, n=5), but no indels at *B2M* and *CIITA* in the *TRAC*-CAR KI (Cas12a) + MHC dKO (BE) conditions (**Figure 6a,b**). The analysis confirmed that indels at *B2M* and *CIITA* found in *TRAC*-CAR KI (Cas9) + MHC dKO (Cas9/BE) induce frameshifts leading to efficient KO of the genes (**Suppl. Figure 4 b,d)**. In addition to indels, base editing was detected in *TRAC*-CAR KI (Cas9) + MHC dKO (BE) treated samples, showing that base editing and DSBs occur during the same intervention (*B2M*: 23.5% ± 8.1%, *CIITA*: 61.8% ± 11.2%, mean ± SD, n=5) (**Figure 6 a,b**). The absence of indels in *TRAC*-CAR KI (Cas12a) + MHC dKO (BE) confirms very low likelihood of DSBs at these sites, suggesting low risk of translocations between BE-target sites and the *TRAC* locus.

**Figure 6:**
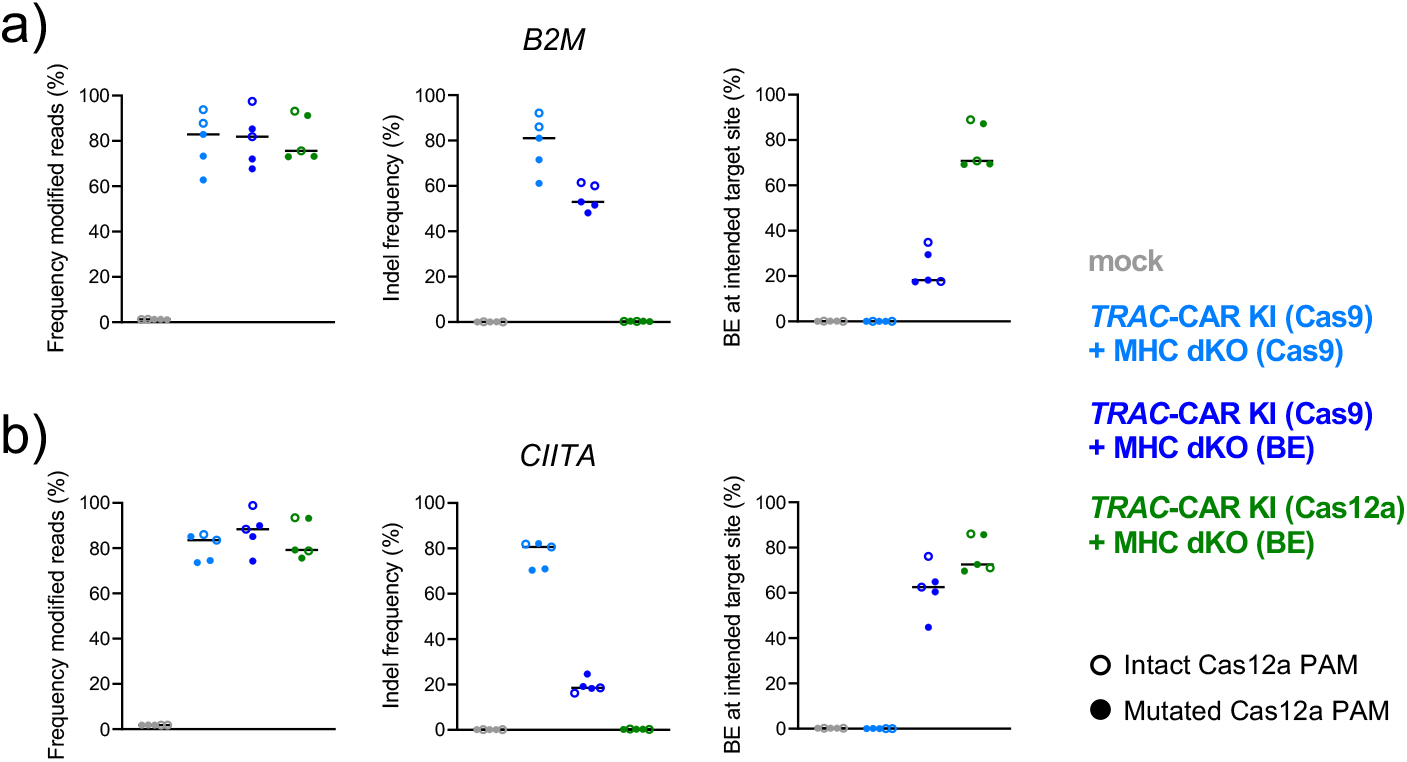
Amplicon sequencing confirms no indel formation at base edited sites when combining Cas12a nuclease and Cas9 BE. Summary of CRIPResso2 analysis showing the frequency of total modified reads, frequency of indels and quantification of intended base editing mediated base changes mapped to *B2M* (a) or to *CIITA* (b). n=5 healthy donors.

### MHC silencing prevents allo-specific T cell cytotoxicity

Higher MHC class I expression in the *TRAC*-CAR KI (Cas12a) + MHC dKO (BE) edited CAR T cells may prevent the intended resistance to allo-specific T cell responses. We generated allo-specific effector T cell lines containing both CD8 and CD4 T cells by repeated restimulation of NK-cell depleted PBMC with irradiated CD3-depleted PBMCs from the CAR T cell donor. After three weeks expansion, we tested the cytotoxicity of the allo-specific T cell lines toward different gene-edited CAR T cell products. As hypothesized, we observed dose-dependent elimination of unmodified donor T cells (mock) and CAR T cells with intact MHC (*TRAC*-CAR KI (Cas12a) + sgRNAs). T cells with *TRAC*-KO (Cas9) + MHC dKO (BE), *TRAC*-CAR KI (Cas9) + MHC dKO (Cas9/BE) and also *TRAC*-CAR KI (Cas12a) + MHC dKO (BE) were not lysed, even at effector to target ratios of 8 : 1 and 12h incubation (**Suppl. Figure 5**). We conclude that the downregulation of MHC class I by *B2M* splice site base editing in combination with complete elimination of MHC class II is sufficient to prevent allo-specific T cell attack.

## IV. Discussion

Multiplex gene editing with a single conventional CRISPR-Cas nuclease system can induce high rates of translocations in cell products (**Figure 1**). Albeit enabling a significant reduction, simultaneous editing with a Cas9 nuclease and a corresponding Cas9 BE led to high rates of translocations through gRNA exchange between the editors (**Figure 2+6**). Combining different CRISPR-Cas systems – here: Cas12a Ultra nuclease and the Cas9 BE ABE8.20m – mitigated gRNA swap and enabled multiplex editing with minimal translocations. In our proof-of-principle model, this completely virus-free solution alleviated the need for multiple, repeated transfections. We thereby demonstrate a simplified strategy for manufacturing of complex gene-edited CAR T cell products that enhances the translational feasibility of CRISPR/Cas edited, allogeneic off-the-shelf applications.

Genotoxicity is a concern for the clinical translation of genetically engineered cell products. In this study, we have quantified balanced translocations as an indicator for chromosomal rearrangements which may affect the safety of multiplex-edited cell products. As our strategy prevents the occurrence of multiple DSBs at different genomic sites, we expect unbalanced translocations, homology-mediated translocations, and translocations with off-target sites to also be reduced. However, unbiased detection methods are necessary to verify the absence of unexpected chromosomal alterations in a potential clinical product. As HDR is still required for the KI of the CAR construct, the necessary DSB leaves an inherent risk for large deletions, homology-mediated translocations and aneuploidy at the on-target site^41,42^. Therefore, safety evaluation of gene-edited cell therapy candidates should also include a comprehensive off-target analysis, which is out of the scope of the current study.

Multiple approaches have been described to reduce or mitigate undesired outcomes during multiplex editing. For instance, promoting HDR at DSBs can reduce the frequency of large deletions^43^ and translocations^21^ induced by CRISPR nucleases. Interestingly, in bulk edited cells, the HDR-mediated TRAC KI did not seem to have a higher abrogating effect on translocations involving the *TRAC* gene, relative to the *B2M* and *CIITA loci* (**Figure 1**). We expect that a pure population of *TRAC*-replaced CAR-expressing cells exhibits reduced translocations involving the *TRAC* locus, due to higher frequency of HDR at the DSB^21^. Another alternative would be the temporal separation of the different gene editing steps with programmable nucleases. Subsequent editing using TALEN transfected every 3 days also allows to reduce translocations to a minimum, but this complicates manufacturing and potentially cell viability due to additional cell handling steps^44^. Future studies should optimize double nicking^45^ with the Cas9 BE to install a DSB for HDR alongside other KO, which should avoid translocations as well, but it may be less efficient than use of a bona-fide nuclease for KI.

Increasing the efficacy and reducing the toxicity of gene editing allows optimized cell yields for therapeutic applications. In line with previous reports^22,28,38,46^, co-transfection of base-modified mRNA encoding for 8^th^ generation adenine BEs achieved highly efficient editing at two sites (**Figure 2, 5, 6**). Most groups reported the combination of retroviral gene transfer and multiplex editing in separate steps^22,26,28^. Diorio *et al* combined cytosine base editing for B2M silencing with a Cas12b nuclease for AAV6-assisted GFP KI but did not investigate translocations in this experiment^22^. To our knowledge, non-viral KI has not been combined with multiplex base editing in a single step. With an average of 30% CAR KI rates using a PAM-mutated template, further optimizations could increase the overall yield for allogeneic cell therapy production. We observed significant DNA-dependent toxicity after electroporation (**Suppl. Figure 3**) which may be reduced by careful titration of the dsDNA template, small molecule drugs to temporarily prevent DNA sensing^37^ or the use of nanoplasmids^47^, circularized ssDNA^48^ or linear ssDNA^49^ as donor templates. AAV6’s high efficacy for KI can be explained by superior delivery of templates to the cells’ nuclei, thereby increasing the probability for HDR due to the high density of alternate repair templates. Non-viral KI may be enhanced by adding truncated Cas target sequences (tCTS) that shuttle donor templates to the nuclei. Adding tCTS has shown to improve KIs with linear dsDNA templates ^50^, plasmids^51^ and most impressively with ssDNA^52^. HDR enhancing drugs could be applied to further increase KI rates^37,52,53^.

Allogeneic T cell products would offer a solution to increase the accessibility of T cell therapies by lowering the price and avoiding logistical challenges of personalized manufacturing. Replacing the TCR can prevent GvHD^35^ but the infused T cells are at risk of rejection by allo-reactive T cells due to MHC mismatches^54^. Thus, the depletion of MHC-I and optionally MHC-II have been exploited to prevent T cell mediated rejection^34^. MHC-I deficient cells could be susceptible for NK killing by “missing self” activation^54^. Therefore, additional genetic modifications may be required to enable optimal long-term persistence of an allogeneic cell product in immunocompetent patients. Overexpression of NK-inhibitory receptors, such as CD47 or the non-polymorphic single-chain HLA-E-B2M fusion molecule,^44^ could be performed. Additionally, deletion of NK-cell activating receptors, such as CD155, has been proposed to protect hypoimmunogenic cells^55^. Furthermore, intensive lymphodepletion with an anti-CD52 antibody has been established to prolong the persistence of a CD52-silenced universal CAR T cell product^14,15,22^.

The diversity of CRISPR-Cas systems can be exploited to achieve highly efficient and complex editing without translocations. In addition to the presented off-the-shelf solution, the same combination of Cas12a and Cas9 BE could be used to create autologous cell products with drug resistance^14,15,56,57^, silenced immune checkpoints^16,23,24^, modifications to avoid fratricide^58,59^ or enhance anti-tumor effects^60,61^. Combining different CRISPR-associated nucleases would also enable complex genetic interventions for other manipulated cell products or even in vivo applications, e.g. HDR for gene corrections combined with epigenetic editing, base editing or RNA editing. Therefore, our study showcases the potential of directing different effector enzymes to distinct sites in the human genome, all within a single intervention.

## V. Methods and Materials

### Ethical statement

The study was performed in accordance with the Declaration of Helsinki. Peripheral blood from healthy human donors was obtained after informed and written consent (Charité ethics committee approval EA4/091/19).

### PBMC isolation and T cell enrichment

PBMCs were isolated by layering the whole blood onto Biocoll separating solution (Bio&SELL, Germany) in 50mL Leucosep Tubes (Greiner, Germany) and using density-gradient centrifugation as previously described^37^. CD3^+^ T cells were positively enriched using magnetic column enrichment with human CD3 microbeads according to the manufacturer’s recommendations (LS columns, Miltenyi Biotec, Germany).

### Cell culture

T cells were cultured in T cell medium containing RPMI 1640 (PAN Biotech), 10% heat-inactivated fetal calf serum (FCS) (Biochrom), recombinant IL-7 (10 ng/mL, Cell-Genix) and IL-15 (5 ng/mL, Cell-Genix). All cell culture experiments were performed at 37°C and 5% CO2. Polyclonal T cell stimulation was performed for 48 h on anti-CD3/anti-CD28-coated tissue culture plates. Coating of vacuum gas plasma-treated polystyrene 24-Well-Tissue-Culture plates (Corning) was performed overnight with 500 μL/well of sterile ddH2O (Ampuwa) supplemented with 1 μg/mL anti-CD3 monoclonal antibody (mAb) (clone OKT3; Invitrogen) and 1 μg/mL anti-CD28 mAb (clone CD28.2; BioLegend). Plates were washed twice in PBS and once in RPMI without letting the wells dry out. T cells were seeded at a density of 1–1.5 × 10^6^ per well in a 24-well-plate. For allogeneic T cell generation, PBMCs were depleted of NK cells using LD columns and the NK cell isolation kit, human (Miltenyi Biotec, Germany). Subsequently, allo-reactive T cells were stimulated 1:1 by adding irradiated CD3-depleted PBMCs from a different donor at the day of isolation. Allo-specific T cells were re-stimulated in a 1:1 ratio with the same CD3-depleted PBMCs on day 5 after isolation and expansion

### Generation of Cas9 base editor mRNA

The ABE8.20-m plasmid was kindly provided by Nicole Gaudelli (Addgene plasmid # 136300)^38^. The adenine base editor ABE8.20-m was cloned into a plasmid containing a dT7 promoter followed by a 5′ untranslated region (UTR), Kozak sequence, the ABE sequence and a 3′ UTR (proprietary backbone plasmid by Trilink). The plasmid was used as a template for PCR (KAPA HiFi HotStart 2x Readymix; Roche) with a forward primer (F: 5’-CGCGGCCGCTAATACGACTCAC-3’) correcting the mutation in the T7 promotor and a reverse primer adding the 120 bp long polyA (R: 5’-TTTTTTTTTTTTTTTTTTTTTTTTTTTTTTTTTTTTTTTTTTTTTTTTTTTTTTTTTTTTTTTTTTTTTTTTTTTTTTTTTTTTTTTTTTTTTTTTTTTTTTTTTTTTTTTTTTTTTTTTCTTCCTACTCAGGCTTTATTCAAAGACCA-3’). The PCR product was purified using DNA Clean & Concentrator-5 kit (Zymo Research), followed by the in vitro transcription using the HiScribe™ T7 High Yield RNA Synthesis Kit (New England Biolabs (NEB)) with N1-methyl-pseudouridine (Trilink) instead of uridine and co-transcriptional capping with CleanCap AG (TriLink Biotechnologies) with 1 μg linear PCR product as a template. To remove template DNA, 70 μL Ambion nuclease-free water (Life Technology Corp.), 10 μL of 10X DNase I Buffer (NEB) and 2 μL of RNase-free DNase I (NEB) were added to 20 μL of IVT reaction and incubate at 37°C for 15 minutes The mRNA was purified using the Monarch® RNA clean up kit (NEB) and the purity was checked using a 1.5 % agarose gel using (2X) RNA Loading dye (NEB) and ssRNA Ladder (NEB) after denaturation at 70°C for 10 min and placing the RNA on ice for 2 min. The mRNA was quantified using the Nanodrop 1000 (Thermo Fisher Scientific) and stored at −80°C prior to use.

### Generation and modification of dsDNA HDRT for insertion of a chimeric antigen receptor

The HDR donor templates were used for targeted insertion of a second generation CD19 CAR based on the original FMC63 single-chain variable fragment (scFv) with an intermediate-length IgG1 hinge, a CD28 transmembrane and costimulatory domain linked to a cytosolic CD3 zeta domain (**Suppl. Table 1**). The HDRTs were generated as previously described^37^ and modified for improved KI with the Cas12a enzyme by mutating the Cas12a PAM. Multiple fragment In-Fusion cloning was performed according to the manufacturer’s protocol (Clontech, Takara) with purified PCR fragments (Kapa Hotstart HiFi Polymerase Readymix, Roche) after using primers introducing the desired change and generating overlaps of 15 bp (**Suppl. Table 1**). In-Fusion cloning strategies were planned with SnapGene (Insightful Science; snapgene.com). In-Fusion reactions were performed in 5 μL reactions at the recommended volume ratios. 2.5 μL of In-Fusion reaction mixtures was transformed into 10 μL of Stellar Competent E. coli, plated on ampicillin containing (LB) broth agar plates and incubated at 37°C overnight. After performing colony PCR for size validation with universal primers adjacent to the pUC19 insertion site (M13-for: 50 - GTAAAACGACGGCCAG-30; M13-rev: 50 -CAGGAAACAGC TATGAC-30), 5 mL ampicillin-containing LB medium was inoculated with the selected clones and incubated at 37°C overnight. Plasmids were purified using ZymoPURE Plasmid Mini Prep Kit (Zymo Research). Sequence validation of HDR-donor-template-containing plasmids was performed by Sanger Sequencing (LGC Genomics, Berlin). The TRAC CD19-CAR HDR template was amplified from the plasmid by PCR using the KAPA HiFi HotStart 2x Readymix (Roche) with a reaction volume of 500 μL and the primers in **Suppl. Table 1**. PCR products were purified and concentrated using paramagnetic beads (AMPure XP, Beckman Coulter Genomics), including two washing steps in 70% ethanol. HDRT concentrations were quantified using the Qubit 4 fluorometer (Thermo Fisher Scientific) and a Qubit™ dsDNA BR-Assay-Kit according to the manufacturer’s protocol and adjusted to 1 μg/mL in nuclease-free water.

### RNP formulation for gene editing

Synthetic modified gRNA (sgRNA: Cas9 or crRNA: Cas12a) sequences which were previously described^38,37,49^ were purchased from Integrated DNA Technologies (IDT), carefully resuspended in nuclease-free 1x TE buffer at 100 μM concentration, aliquoted and stored at −20°C prior to use (**Suppl. Table 2**). Per electroporation of 1-1.5×10^6^ primary human T cells, 0.5 μL of an aqueous solution of 15- to 50-kDa poly(L-glutamic acid) (PGA) (Sigma-Aldrich, 100 μg/μL) was mixed with 0.48 μL of *TRAC*-specific modified sgRNA (Cas9) or *TRAC*-specific modified crRNA (Cas12a) by pipetting thoroughly. Then, 0.4 μL recombinant *Streptococcus pyogenes* Cas9 protein (Alt-R S.p. Cas9 Nuclease V3; IDT; 10 μg/μL = 61 μM) or *Acidaminococcus sp. BV3L6* Cas12a (Alt-R A.s.Cas12a (Cpf1) Ultra; IDT; 10 μg/μL = 63 μM) are added and mixed by thorough pipetting. The molar ratio of Cas9/Cas12a and sgRNA was ∼ 1:2. The mixture was incubated for 15 min at room temperature (RT) to allow for RNP formation and placed on ice. For KI experiments, 0.5 μL HDRT (1 μg/μL) per 10^6^ cells was added at least 5 minutes prior transfection.

### Transfection of gene editors

Primary human T cells were harvested approx. 48 hours after anti-CD3/CD28 stimulation and washed twice in sterile PBS by centrifugation with 100x*g* for 10 min at RT. Depending on the condition, modified mRNA (2 μg, unless stated otherwise) and/or additional sgRNA (0.48 μL each) was added to 1.88 μL of RNP/HDRT suspension. The harvested cells were resuspended in 20 μL ice-cold P3 electroporation buffer (Lonza) for electroporation of 1-1.5 × 10^6^ cells. The exposure time to the electroporation buffers was kept to a minimum, 20 μL of resuspended cells were transferred to the RNP/HDRT (+mRNA and sgRNA) suspension, mixed thoroughly and transferred into a 16-well electroporation strip (20 μL = 1-1.5 10^6^ cells per well, Lonza). Prior to electroporation, the strips or cartridges were gently tapped onto the bench several times to ensure the placement of the liquid on the bottom of the electroporation vessel without any trapped air (bubbles). Electroporation was performed on a 4D-Nucleofector Device (Lonza) using the program EH-115. Directly after electroporation, pre-warmed T cell medium was added to the cells (90 μL per well). Afterward, the cells were carefully resuspended and transferred to 96-well round-bottom plates (50 μL/well) containing 150 μL pre-warmed T cell medium per well at a density of 0.5 × 10^6^ cells per well.

### T cell expansion after electroporation

First medium change or first splitting of cells was performed 18 h after electroporation, unless stated otherwise. Cells were expanded in T cell medium on 96-well round-bottom plates or 24-well plates. T cells were split when culture-medium turned orange/yellow, indicating pH-change. Typically, within the first two to three days after electroporation, T cells were split (50:50) every day or every other day. Later T cells were split, or medium was changed every two to three days. Depending on the readout, some of the T cells were pelleted and stored at −20°C until genomic DNA extraction. In other experiments, T cells were further expanded, counted on day 1, 4, 7 and 14 using flow cytometry to track the expansion, followed by cryopreservation in freezing medium (FCS containing 10% DMSO).

### Flow cytometry

Flow cytometry analysis was used to determine the number of viable cells and the gene editing efficiency on a protein level. Measurements were performed on a Cytoflex LX device (Beckman Coulter Genomics) using 96-well flat-bottom plates for cell counts and 96-well U-bottom plates for other measurements. For cell counting, 10 μL of cells were diluted 1:10 in PBS with DAPI (1:10000) before acquiring 20 μL. For determination of the editing efficiency, approximately 100,000 T cells were transferred onto the 96-U-bottom-well plate and washed by adding 200 μL of PBS, centrifuging the plates at 400g for 5 min at RT, discarding the supernatants, and resuspending the pellets in the remaining volume by vortexing briefly. For any individual staining procedure, a mastermix of the fluorophore conjugated antibodies diluted in PBS was prepared. 20 μL of the mastermix was added per well. The plates were incubated for 15 min at 4°C. Due to potential cross-reactivity of the anti-Fc antibody, which we used for CAR staining, a first extracellular staining step with anti-Fc antibody and a live-dead discriminating dye followed by two washing steps was performed prior to staining with anti-CD3 PacBlue, anti-HLA-A,B,C PE-Cy7 and anti-HLA-DR,DQ,DP FITC.

| Name | Antigen | Dye | Dilution | Host Species | Reactivity | Clone | Catalog number | Manufacturer |
| --- | --- | --- | --- | --- | --- | --- | --- | --- |
| LIVE/DEAD Fixable Blue | - | - | 1:100 | - | - | - | L23105 | Invitrogen |
| Fc AF647 | IgG1, Fc-gamma fragment | AF647 | 1:50 | goat | human | polyclonal | 109-605-098 | Jackson Immuno Research |
| CD3 PacBlue | CD3 | Pacific Blue | 1:50 | mouse | human | UCHT1 | A93687 | Beckman Coulter |
| HLA-A,B,C PE Cy7 | HLA-A,B,C | PE-Cy7 | 1:100 | mouse | human | W6/32 | 311429 | Biolegend |
| HLA-DR,DQ,DP FITC | HLA-DR,DQ,DP | FITC | 1:100 | mouse | human | Tü39 | 361705 | Biolegend |

### Cytotoxicity assay of allo-reactive T cells against multiplexed edited T cells

Leftover CD3 negative cells obtained from CD3 enrichment for the generation of gene-edited T cells, were used to stimulate and enrich allo-reactive T cells from a different donor. The expanded alloreactive T cells were co-cultured with mock electroporated, or gene-edited target cells with disrupted HLA-I and -II genes. The target cells were fluorescently labeled with CFSE (Thermo Fischer Scientific). Allo-reactive T cells were added to 25,000 target T cells in 96-well, round-bottom, cell-culture plates at seven different effector:target cell ratios (8:1, 4:1, 2:1, 1:1, 0.5:1, 0.25:1, 0.125:1) with control wells containing only the target cells. The plates were centrifuged at 100g for 3 min at RT and incubated at 37°C and 5% CO_2_. After 4 and 16h, 80 μL of resuspended cell suspension was added to 80 μL of PBS containing 1:10,000 DAPI followed by a minimum of 10 min incubation at 4°C. 30 μL were analyzed by flow cytometry. Alloreactive T cell mediated cytotoxicity was calculated by the reduction in cell number of CFSE labeled target cells in the co-culture, compared to the target only control. The experiment was performed on day 23 after blood collection (day 21 after electroporation).

### Digital droplet polymerase chain reaction (ddPCR) for translocation quantification

ddPCR assays were designed for balanced translocations between the TRAC, B2M or CIITA gene (**Suppl. Table 3**). A ddPCR assay for the RPP30 gene was used as reference^62^. 50 ng of HindIII-HF (NEB, Germany) digested gDNA were used as template for a 20 μL PCR reaction with 1 μl (5μM) of the forward and reverse primers for both the target and reference genes, 1 μL (5μM) of target and reference probe, 10μL of 2X ddPCR Supermix for Probes (No dUTP) and nuclease free water. Droplet generation was performed with the QX200 Droplet Generator, using 20 μL of sample and 70 μL Droplet Generation Oil inside a DG8 Cartridge covered by a DG8™ Gaskets for QX200™/QX100™ Droplet Generator. Any empty wells were filled with ddPCR Buffer Control. After droplet generation, droplets were gently pipetted into a 96-well PCR plate and sealed using the PX1™ PCR Plate Sealer and pierceable foil heat seal for 5s at 185°C. Samples with less than 15,000 accepted droplets or more than 99% positive droplets for the reference gene were discarded and repeated.Primers and probes were ordered from IDT and all other devices and consumables were acquired from BioRad.

### Optimization of ddPCR assays to reduce background signal by HDRT

For balanced translocations between *TRAC* and *B2M/CIITA*, we initially observed a background signal for ddPCR assays with a forward primer binding to the TRAC homology arm of the HDRT, leading to false-positive events in ddPCR four days after transfection (TRAC_B2M, **Suppl. Fig. 5a,b,c**). When the T cells were expanded until d14, the noise disappeared, presumably due to degradation of the HDRT (**Suppl. Fig. 6 a,e,f**). When the reporter probe was placed on the *B2M* instead of the *TRAC* locus, the background was eliminated (**Suppl. Fig. 6 b,g,h**). Overall, at day 4 after transfection, no significant difference was observed in the total translocation frequency between unedited (0.05%, SD=0.06) and *TRAC*-CAR KI (Cas12a) + MHC dKO (BE) samples (**Figure 6f**). The specificity of the assays were validated by PCR of genomic DNA from *TRAC*-KO (Cas9) + MHC dKO (Cas9) (triple KO) samples and synthetic gene fragments (gBlocks™ (gB) by Integrated DNA Technologies Inc) modeling the translocations as positive controls (**Suppl. Fig. 6i**) The test PCRs were performed using Red Taq DNA Polymerase Master Mix (VWR) and the GeneRuler™ 100 bp Plus DNA Ladder (Thermo Fisher Scientific).

### Sanger sequencing - editR

Genomic DNA was isolated on day 4 after electroporation using the Quick-DNA Miniprep plus Kit (Zymo Research). Primers were designed to PCR amplify a 500-700 bp fragment of the *B2M* and *CIITA* locus from gDNA using Cosmid ^63^. Specificity of the primers was checked *in silico* using primer-Blast^64^ and on a 1.5% agarose gel. The fragment was purified using the DNA Clean & Concentrator-5 kit (Zymo Research). Sanger sequencing was performed (LGC Genomics) and the base editing efficiency was quantified using EditR in RStudio^65^.

### Amplicon sequencing

Genomic DNA was isolated on day 4 after electroporation unless otherwise stated. 200 bp PCR amplicons (KAPA HiFi HotStart 2x Readymix; Roche) of the *B2M* and *CIITA* loci containing DNA adapters were generated using primer pairs and adapters published by Gaudelli et al ^38^. The PCR products were purified using AMPure beads (AMPure XP, Beckman Coulter Genomics) and the size was checked on a 1.5% agarose gel. In a second PCR (KAPA HiFi HotStart 2x Readymix; Roche), dual indices were added using primers binding to the previously attached adapters (**Suppl. table 4**). The PCR products were purified using AMPure beads (AMPure XP, Beckman Coulter Genomics) and the size was checked on a 1.5% agarose gel. The concentration was measured using the Qubit 4 Fluorometer (Thermo Fisher Scientific) and adjusted to 4 μM. 20 μL (4 μM) of each samples were united to a library and diluted to a 1 μM concentration. 5 μL of Library (1 μM) were denatured using 5 μL of 0.1 N NaOH (Sigma) and diluted to a loading concentration of 1.4 pM. The library was spiked with 20% of diluted and denatured 1.4 pM PhiX control (Illumina) and 500 μL of the final sample were sequenced 2×150 cycles using the MiniSeq Mid Output Kit (300-cycles) (Illumina) on an Illumina MiniSeq instrument.

### Data analysis, statistics, and presentation

NGS data was demultiplexed using local run manager (Illumina). For base editing efficiency and Indel quantification CRISPresso2^66^ was run in batch mode for the *B2M* and *CIITA* amplicons using the following settings: 1) BE: -p 4 --base_edit -wc -10 -w 10 --min_average_read_quality 30 --conversion_nuc_from A --conversion_nuc_to G 2) NHEJ: --p 4 -min_average_read_quality 30. The data of all samples was pooled and the frequencies of reads with the intended base change were calculated using python (**Suppl. Note 1**) (Python Software Foundation, version: 3.7.14), Available at http://www.python.org). Flow cytometry data were analyzed with FlowJo software v.10 (BD Biosciences). Data from different assays were collected in Excel (Microsoft). Graphs and statistical analyses were created using Prism 9 (GraphPad). Conditions with failed electroporation (indicated by 4D-Nucleofector Device) were recorded during the experiment and excluded from analysis. Schemes and graphs in the presented figures were created using www.biorender.com.

## VI. Data availability statement

The ABE8.20m was a gift from Nicole Gaudelli (Addgene plasmid #136300). The IVT plasmid backbone contains proprietary 5’-UTR and 3’-UTR sequences and was procured from TriLink Inc. under a non-disclosure agreement. All other construct sequences can be found in **Suppl. Table 5**. All other data can be obtained from the corresponding author upon request.

## VII. Acknowledgements

We would like to thank Nicole Gaudelli (Beam Tx) and Jason Gehrke (Beam Tx) for sharing the ABE8.20-m plasmid and information on protocols for base-modified mRNA production. Alexej Knaus (University of Bonn) and Daniel Ibrahim (MPI/BCRT) for help in the planning of the NGS experiments. Jaspal Kaeda (Charité), Heiko Trautmann (Kiel University) and Marco Lodrini (Charité) for theoretical input and technical advice on the design of translocation assays and ddPCRs.

## VIII. Funding

This project has received funding from the European Union’s Horizon 2020 research and innovation program under grant agreement no. 825392 (ReSHAPE-h2020.eu) to H-DV, MS-H, PR, DLW, and the SPARK-BIH program from the Berlin Institute of Health to JK and DLW.

## IX. Author contributions

VG designed the study, planned and performed experiments, analyzed results, interpreted the data, and wrote the manuscript. CFlugel, JK, WD performed experiments, analyzed results, interpreted data, and edited the manuscript. VD, CFranke, MS performed experiments and analyzed results. AP provided reagents. H-DV, MS-H, PR provided reagents, interpreted data, and edited the manuscript. DLW designed and led the study, planned experiments, interpreted data, and wrote the manuscript. All authors discussed, commented on, and approved the manuscript in its final form.

## X. Conflict of interest statement

Charité has received reagents for gene editing (TRAC sgRNA, custom gblocks) from Integrated DNA technologies Inc as part of a collaboration agreement.

## Supplementary Figures

**Supplementary Figure 1:**
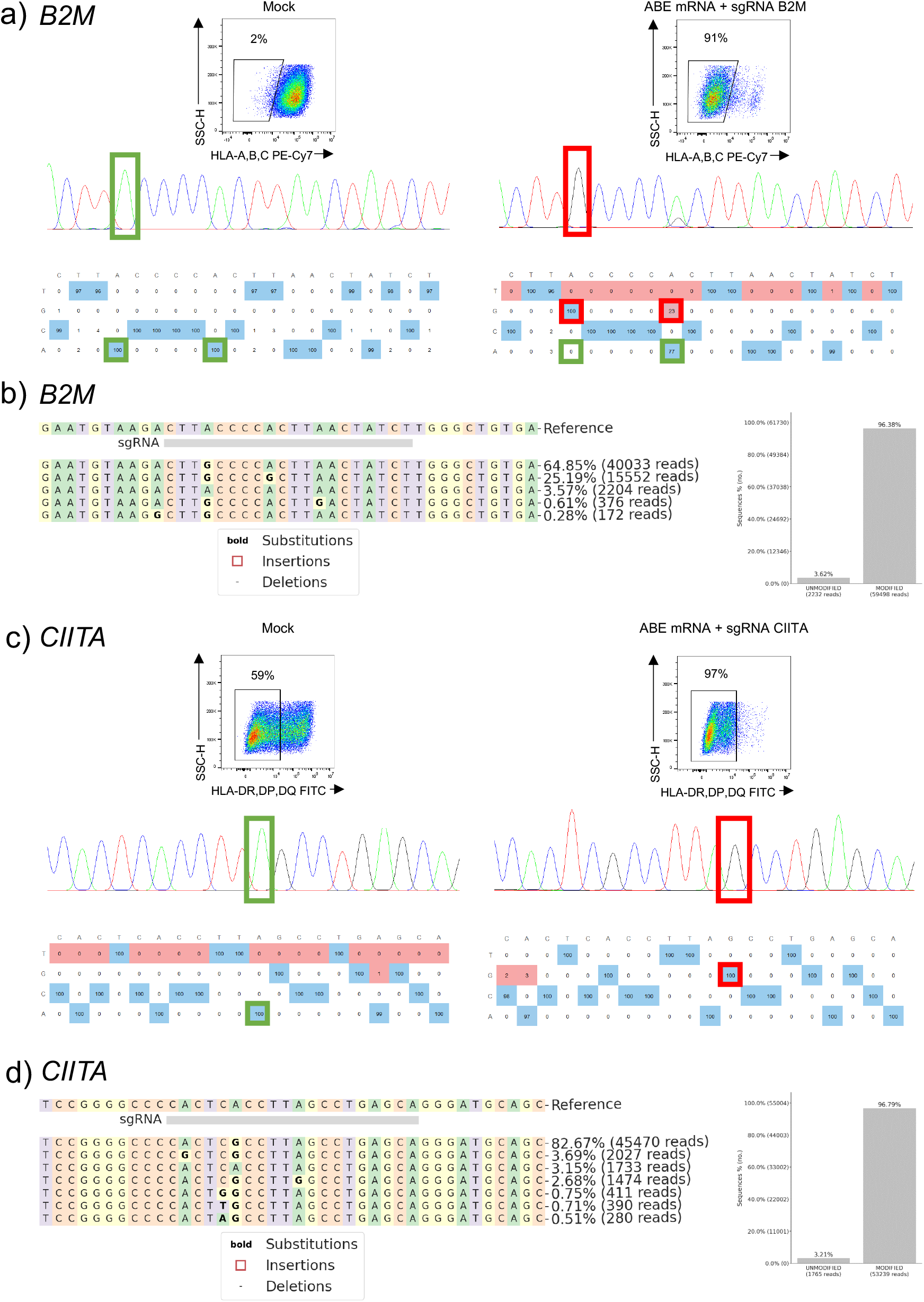
Sequencing confirms efficient base editing at *B2M* and *CIITA* after transfection of Cas9 adenine base editor mRNA. T cells were mock-electroporated or transfected with mRNA encoding for the adenine base editor ABE8.20-m with a gRNA targeting a splice site of *B2M* or *CIITA*. Analysis was performed 4 days after transfection. (a+c) Representative flow cytometry and EditR results from Sanger sequencing data at *B2M* locus (a) or *CIITA* locus (c). b+d) Representative results from targeted next generation sequencing confirms high rate of A to G conversions in modified, but not unmodified cells (left: top 5 reads, right: summary of modified reads according to CRISPResso2 analysis) at *B2M* (b) and *CIITA* (d) locus). n=3 healthy donors.

**Supplementary Figure 2:**
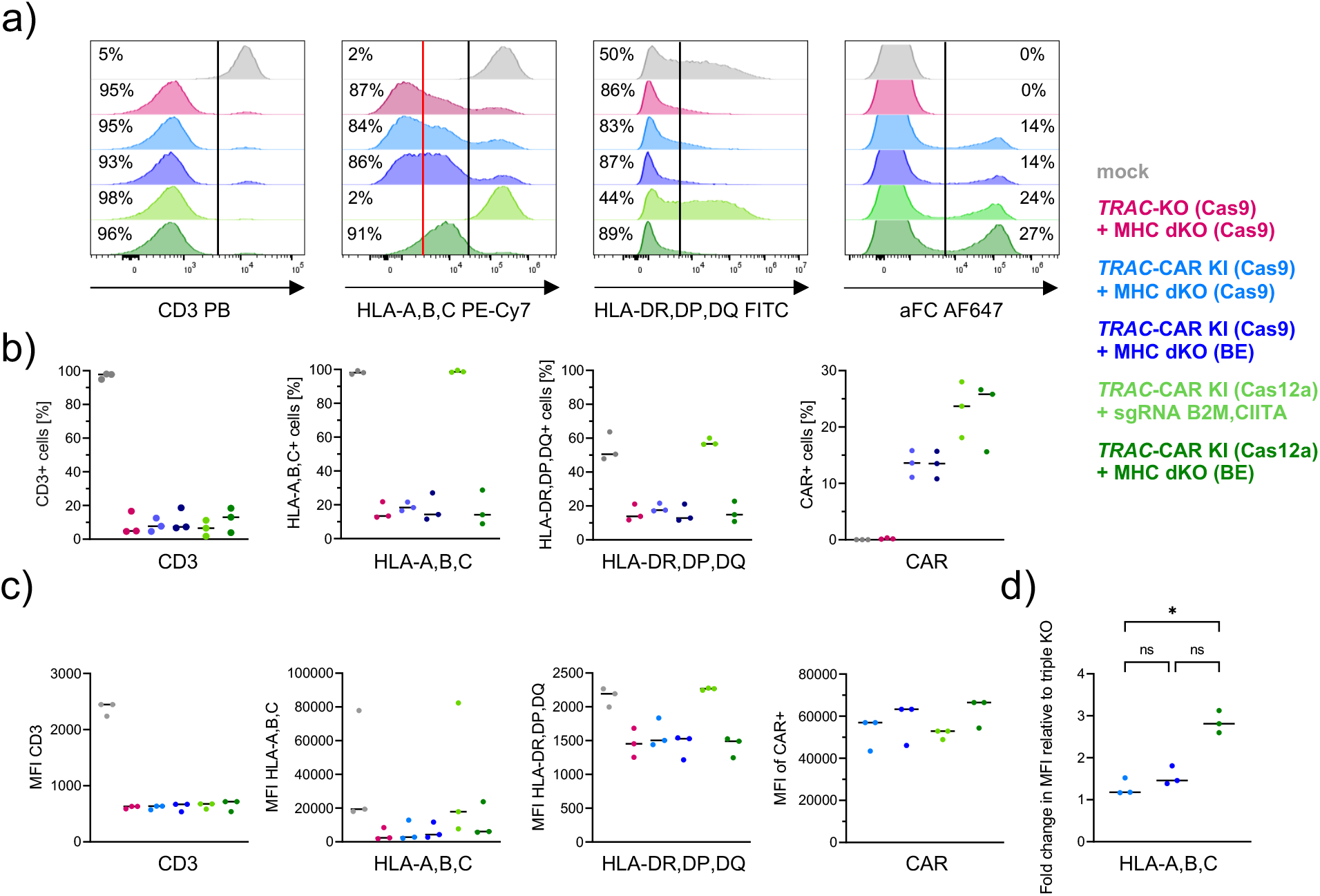
Comparison of surface expression of CAR, TCR and MHC expression in different multiplex edited T cells. a) Representative flow cytometry histograms for surface expression of CD3, HLA-A,B,C, HLA-DR,DP,DQ or CAR (aFC) four days after T cells were transfected with or without gene editors. b) Paired summary data for surface marker expression from T cells of three independent donors four days after treatment. c) Summary data of median fluorescent intensity (MFI) for the respective markers after gene editing. d) Normalized MFI data comparing the relative expression level of edited fraction (left of black line in histograms in a) of *TRAC*-KO (Cas9) + MHC dKO (Cas9) cells with different gene edited T cell populations. n=3 independent healthy donors. Statistical analysis of flow cytometry and ddPCR data from 3 donors was performed using a one-way ANOVA of matched data with Geisser-Greenhouse correction. Multiple comparisons were performed by comparing the mean of each column with the mean of every other column and corrected by the Turkey test. Asterisks represent different p-values calculated in the respective statistical tests (ns : p ≥ 0.05; * : p < 0.05; ** : p < 0.01; *** : p < 0.001; **** : p < 0.0001)

**Supplementary Figure 3:**
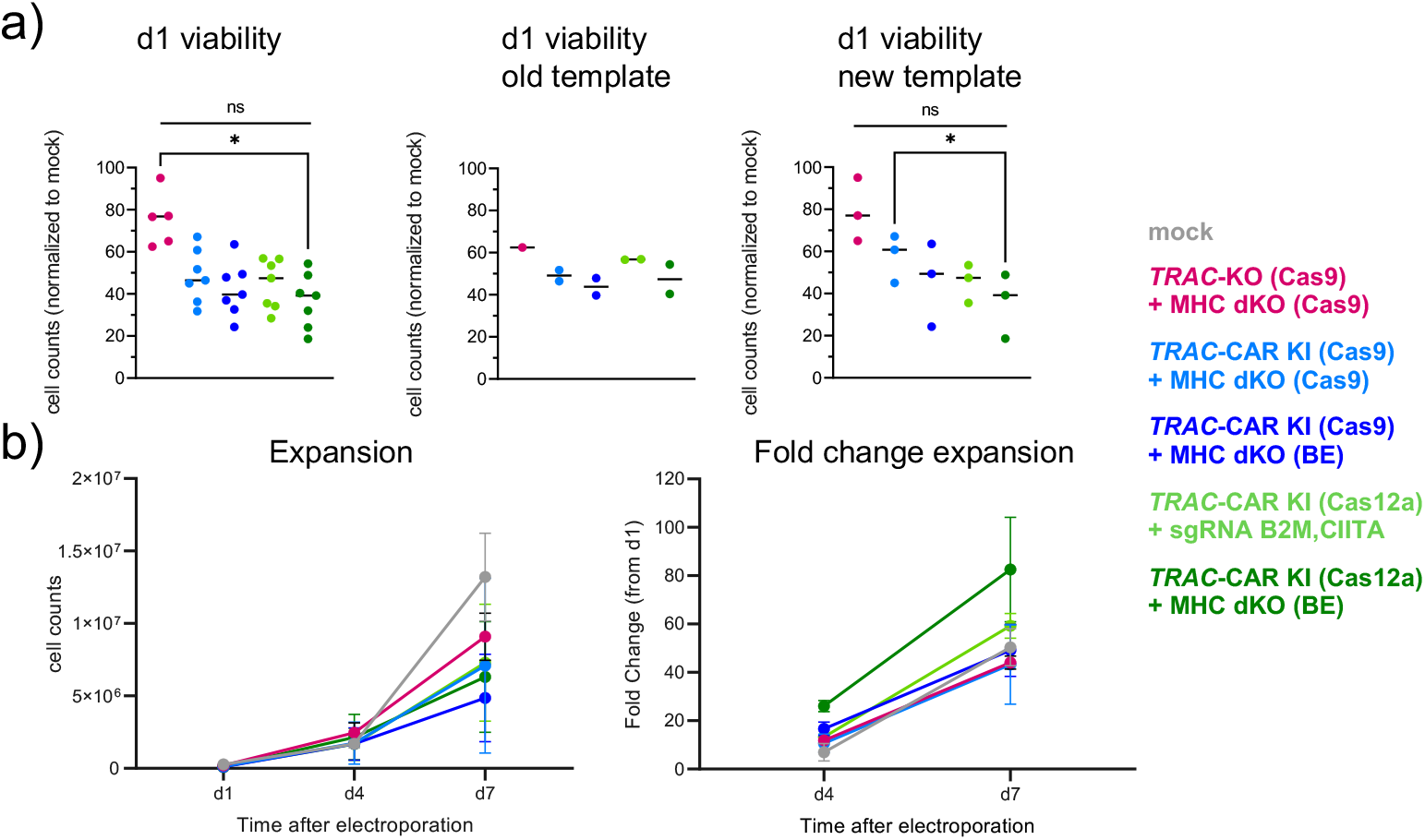
Co-delivery of modified mRNA for base editing does not reduce T cell viability and expansion capacity during non-viral knock-in. a) Summary results of cell viability determined 24 hours after transfection and normalized to mock-electroporated T cells. Followed by individual viability results depending on which HDRT template was used (new: PAM-mutated template as displayed in Figure 3). b) Total cell counts within 7 days after nucleofection and summary of fold expansion for different gene-edited T cells normalized to cell counts from 24 hours after transfection (n=2). Statistical analysis of flow cytometry and ddPCR data from 3 donors was performed using a one-way ANOVA of matched data with Geisser-Greenhouse correction. Multiple comparisons were performed by comparing the mean of each column with the mean of every other column and corrected by the Turkey test. Asterisks represent different p-values calculated in the respective statistical tests (ns : p ≥ 0.05; * : p < 0.05; ** : p < 0.01; *** : p < 0.001; **** : p < 0.0001)

**Supplementary Figure 4:**
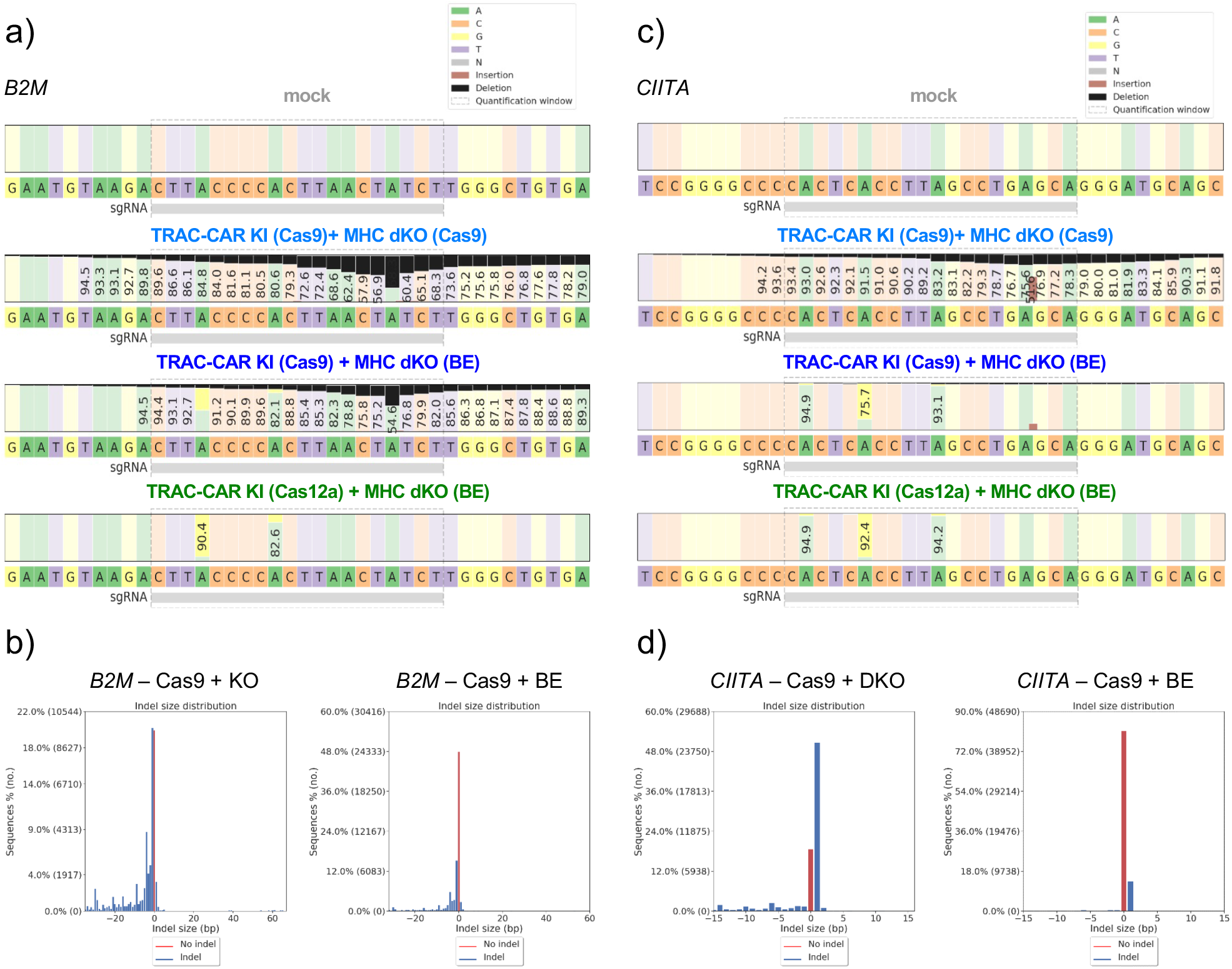
CRISPResso2 results after amplicon sequencing of *CIITA and B2M*. Representative CRISPResso2 results from amplicon sequencing data and Indel size distribution of the a),b) B2M and c,d) CIITA target site from T cells electroporated with different gene editors as described before. n=5 healthy donors.

**Supplementary Figure 5:**
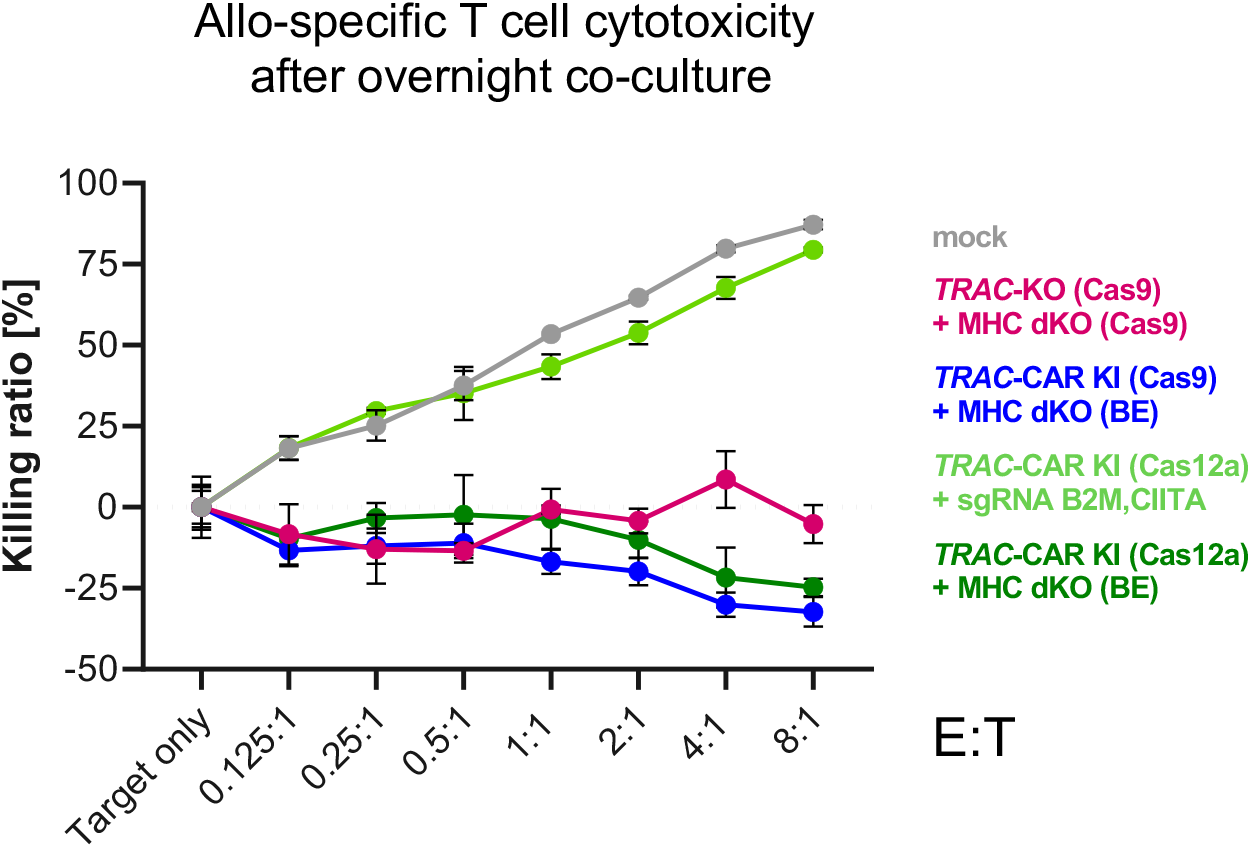
MHC silencing prevent allo-specific T cell cytotoxicity. Allo-specific T cells were generated by stimulating NK cell-depleted PBMCs from donor A with irradiated T cell-depleted PBMCs from the donor used for CAR T cell generation. Allo-specific T cells were re-stimulated twice with irradiated T cell-depleted PBMCs from the CAR T cell donor at day 7 and 14 after isolation. Then, allo-specific T cells were used as effector cells during an overnight killing assay. To this end, CAR T cells were labeled with CFSE. Co-cultures were set up at different effector (allo-specific T cells) to target (edited T cells) ratios and incubated overnight prior flow cytometry analysis. Absolute counts for CFSE-negative T cells were quantified. Results were normalized to target only control. Thick lines indicated mean value, error bars indicate standard deviation. n=3.

**Supplementary Figure 6:**
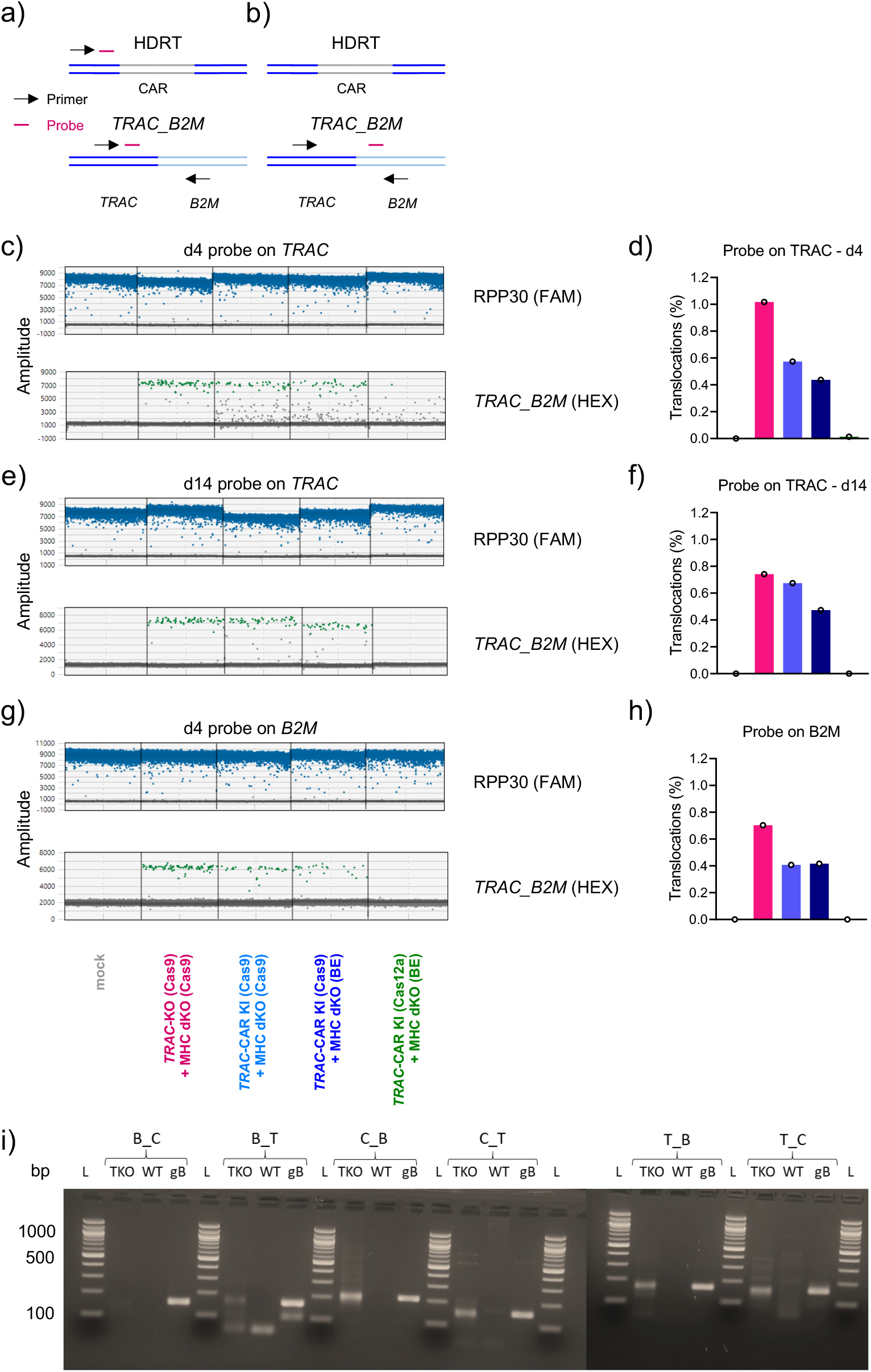
High background in ddPCR assay with probe binding TRAC HDRT removed by placing the probe on non-TRAC locus of translocation. a) ddPCR assay to detect *TRAC_B2M* translocations with the probe binding to the genomic *TRAC* locus as well as to the homology arm of the HDRT. b) Probe binding to B2M prevents binding to HDRT in ddPCR assay detecting *TRAC_B2M* translocations. c) ddPCR raw data using assay described in a) 4 days after gene-editing intervention and d) analysis of different conditions. e) ddPCR raw data using assay described in a) 14 days after gene-editing intervention and f) analysis of different conditions. g) ddPCR raw data using assay described in b) 4 days after gene-editing intervention and h) analysis of different conditions. i) Test PCRs for different ddPCR assays with gene Blocks (gBlocks™, gB) (IDT) as positive control and triple KO cells (TKO).

## Notes

### Competing Interest Statement

Charite has received reagents for gene editing (TRAC sgRNA, custom gblocks) from Integrated DNA technologies Inc as part of a collaboration agreement.

