## Supplemental Note 1 for "Combining different CRISPR nucleases for simultaneous knock-in and base editing prevents translocations in multiplex-edited CAR T cells"

**Supplementary Note 1: Python code for extraction of results after CRISPResso2 batch analysis**

#create new file: results.csv
import csv
with open('results.csv', 'w', newline='') as file:
 wr = csv.writer(file)
 wr.writerow([''])

import os, glob
\# assign directory with CRISPResso2 results
directory = r'C:\Users\...' #insert directory

reference_sequence = "GAATGTAAGACTTACCCCACTTAACTATCTTGGGCTGTGA"

#open Lists to collect results from all files
summary = ['sample name', 'total reads', 'reads after preprocessing', 'reads aligned', 'modified (BE setting)', 'unmodified','WT read frequency', 'BE read frequency', 'undefined Indel or sequence change frequency', 'sum of frequencies']
sample_name = []
total_reads = []
reads_after_preprocessing = []
reads_aligned = []
modified = []
unmodified = []
WT_read_frequency = []
BE_read_frequency = []
undefined_Indel_frequency = []
sum_of_frequencies = []

\# iterate over txt files in that directory and convert to csv
os.chdir(directory)
for file in glob.glob("*.txt"):
 #convert txt to csv
 import pandas as pd
 read_file = pd.read_csv (file, delimiter = '\t')
 read_file.to_csv (file+'.csv', index=None)

\# iterate over Alleles_frequency csv files to get A->G base change information
for file in glob.glob("*Alleles_frequency.txt.csv"):
 #create short sample name
 name = str(file[0:-26])
 print(name)

 #convert csv to dict key=Aligned_Sequence value=percent of reads
 dict_sequence_reads = {}
 List_sample_reads = []
 List_sample_frequencies = []
 colnames = ['Aligned_Sequence', 'Reference_Sequence', 'Unedited', 'n_deleted', 'n_inserted', 'n_mutated', '#Reads', '%Reads']
 data = pd.read_csv(file, names=colnames)
 data.rename(columns={'%Reads': 'freqReads'}, inplace=True)
 \# filter out duplicate reads and add up frequency of same reads
 List_sample_reads = data.Aligned_Sequence.tolist()
 del List_sample_reads[0]
 List_sample_frequencies = data.freqReads.tolist()
 del List_sample_frequencies[0]
 for key,value in zip(List_sample_reads,List_sample_frequencies):
 if key not in dict_sequence_reads:
 dict_sequence_reads[key]=float(value)
 else:
 dict_sequence_reads[key]+=float(value)

 #iterate through Aligned_Sequence in dict
 #create list of BE reads with frequencies and frequency of WT reads
 read_frequencies = 0
 sequences = 0
 sequences_with_BE = 0
 reads_of_WT = 0
 undefined_reads = 0
 List_of_BE_reads = []
 List_of_BE_reads_frequency = []

 \# Sum up frequency of reads with adenine base editing at 12th base position from NGG PAM
 for sequence in dict_sequence_reads:
 sequences += 1
 if sequence[13] == 'G':
 sequences_with_BE += 1
 List_of_BE_reads.append(sequence)
 List_of_BE_reads_frequency.append(str(dict_sequence_reads[sequence]))
 read_frequencies += float (dict_sequence_reads[sequence])
 elif sequence in reference_sequence:
 reads_of_WT += float (dict_sequence_reads[sequence])
 else:
 undefined_reads += float (dict_sequence_reads[sequence])
 frequency_of_all_reads = read_frequencies + reads_of_WT + undefined_reads

 #append results to summary lists
 sample_name.append(name)
 WT_read_frequency.append(reads_of_WT)
 BE_read_frequency.append(read_frequencies)
 undefined_Indel_frequency.append(undefined_reads)
 sum_of_frequencies.append(frequency_of_all_reads)

 #merge list_of_BE_reads and List_of_BE_reads_frequency to csv
 import csv
 with open('results.csv', 'a', newline='') as file:
 wr = csv.writer(file)
 wr.writerow([''])
 wr.writerow(['Found BE reads in sample: ' + name])
 header = ['BE_reads', 'BE_reads_frequency']
 wr.writerow(header)
 for word in list(zip(List_of_BE_reads, List_of_BE_reads_frequency)):
 wr.writerows([word])
 file.close()

#Iterate over csv files of mapping statistics and collect 'READS IN INPUTS', 'READS AFTER PREPROCESSING', 'READS ALIGNED' for summary file
for file in glob.glob("*mapping_statistics.txt.csv"):
 List_total_reads = []
 List_reads_after_preprocessing = []
 List_reads_aligned = []
 colnames_mapping = ['READS IN INPUTS', 'READS AFTER PREPROCESSING', 'READS ALIGNED', 'N_COMPUTED_ALN', 'N_CACHED_ALN', 'N_COMPUTED_NOTALN', 'N_CACHED_NOTALN']
 data = pd.read_csv(file, names=colnames_mapping)
 data.rename(columns={'READS IN INPUTS': 'READS_IN_INPUTS', 'READS AFTER PREPROCESSING':'READS_AFTER_PREPROCESSING', 'READS ALIGNED':'READS_ALIGNED'}, inplace=True)
 List_total_reads = data.READS_IN_INPUTS.tolist()
 del List_total_reads[0]
 List_reads_after_preprocessing = data.READS_AFTER_PREPROCESSING.tolist()
 del List_reads_after_preprocessing[0]
 List_reads_aligned = data.READS_ALIGNED.tolist()
 del List_reads_aligned[0]
 total_reads.extend(List_total_reads)
 reads_after_preprocessing.extend(List_reads_after_preprocessing)
 reads_aligned.extend(List_reads_aligned)

#Iterate over csv files of editing_frequency and collect modified and unmodified reads frequencies for summary file
for file in glob.glob("*editing_frequency.txt.csv"):
 List_modified = []
 List_unmodified = []
 colnames_mapping = ['Amplicon', 'Unmodified%', 'Modified%', 'Reads_in_input', 'Reads_aligned_all_amplicons', 'Reads_aligned', 'Unmodified', 'Modified', 'Discarded', 'Insertions', 'Deletions', 'Substitutions', 'Only Insertions', 'Only Deletions', 'Only Substitutions', 'Insertions and Deletions', 'Insertions and Substitutions', 'Deletions and Substitutions', 'Insertions Deletions and Substitutions']
 data = pd.read_csv(file, names=colnames_mapping)
 data.rename(columns={'Unmodified%': 'freqUnmodified', 'Modified%':'freqModified'}, inplace=True)
 List_unmodified = data.freqUnmodified.tolist()
 del List_unmodified[0]
 List_modified = data.freqModified.tolist()
 del List_modified[0]
 unmodified.extend(List_unmodified)
 modified.extend(List_modified)

#append summary Lists to results.csv
import csv
with open('results.csv', 'a', newline='') as file:
 wr = csv.writer(file)
 wr.writerow([''])
 wr.writerow(summary)
 for word in list(zip(sample_name, total_reads, reads_after_preprocessing, reads_aligned, modified, unmodified, WT_read_frequency, BE_read_frequency, undefined_Indel_frequency, sum_of_frequencies)):
 wr.writerows([word])
 file.close()

**Supplementary Note 1 Python code for extraction of results after CRISPResso2 batch analysis.** Collection of BE read frequency, frequency of total modified reads as well as read frequencies modified by NHEJ (indels) from results of CRISPResso2 batch analysis (BE and NHEJ setting). Representative script for B2M editing is shown. The analysis for CIITA analysis was run after updating the directory, reference sequence and the position of intended base change.
